# Individual differences in the effects of priors on perception: a multi-paradigm approach

**DOI:** 10.1101/523324

**Authors:** Kadi Tulver, Jaan Aru, Renate Rutiku, Talis Bachmann

**Author notes:** corresponding author: Kadi Tulver.

## Abstract

The present study investigated individual differences in how much subjects rely on prior information, such as expectations or knowledge, when faced with perceptual ambiguity. The behavioural performance of forty-four participants was measured on four different visual paradigms (Mooney face recognition, illusory contours, blur detection and representational momentum) in which priors have been shown to affect perception. In addition, questionnaires were used to measure autistic and schizotypal traits in the non-clinical population. We hypothesized that someone who in the face of ambiguous or noisy perceptual input relies heavily on priors, would exhibit this tendency across a variety of tasks. This general pattern would then be reflected in high pairwise correlations between the behavioural measures and an emerging common factor. On the contrary, our results imply that there is no single factor that explains the individual differences present in the aforementioned tasks, as further evidenced by the overall lack of robust correlations between the separate paradigms. Instead, a two-factor structure reflecting differences in the hierarchy of perceptual processing was the best fit for explaining the individual variance in these tasks. This lends support to the notion that mechanisms underlying the effects of priors likely originate from several independent sources and that it is important to consider the role of specific tasks and stimuli more carefully when reporting effects of priors on perception.

## 1. Introduction

### 1.1. Inter-individual differences in perception

Individual differences have been thoroughly researched in several domains of cognitive science, such as intelligence and memory, and have all but dominated the field of personality psychology. In the study of perception, on the other hand, inter-individual variability has traditionally been treated as a source of noise and discarded in favour of studying effects common to groups of people (Calhoun et al., 2001; Kanai & Rees, 2011; Matin, Boff, Kaufman & Thomas, 1986; Rahnev & Denison, 2018; Ulehla, 1966; Wade & Swanston, 2001). In recent years, however, there has been more emphasis on the study of individual differences and specific factors present in perceptual processing, which can offer valuable insight for understanding the cognitive mechanisms underlying perception and behaviour.

While it stands to reason that people who perform better in one perceptual task should also fare well in other similar tasks, the quest for general factors of visual performance has so far yielded mixed results. For example, Verhallen et al (2017) found significant correlations between four paradigms employing face processing and recognition, and proposed the term *f* for the emerging factor that underlies this pattern of positive correlations in the domain of face perception. Other studies have attempted to find a factor for susceptibility to optical illusions (Grzeczkowski et al, 2017; Thurstone, 1944). Unlike Thurstone (1944) who concluded that susceptibility to geometric illusions is one of the 11 factors of visual ability, Grzeczkowski and colleagues (2017) reported that most correlations between their six chosen illusion paradigms were not significant and there was no single factor underlying the performances on these tasks. Likewise, a study by Chamberlain and colleagues (2017) found no substantial evidence to support a monolithic factor structure of local-global processing, although distinct processing biases specific to local and global precedence had previously been suggested (e.g. Milne & Szczerbinski, 2009).

Taken together, more research is required to better understand the structure and qualitative nomenclature of individual differences in visual ability and perception. In the present work we explore the possibility that some systematic individual differences in perception are explained by varying degrees of reliance on prior information.

### 1.2. The role of priors in perception

There is an ever-increasing number of studies challenging one of the most influential views of perception as a process of stimulus-driven bottom-up feature detection and integration (for reviews, see Gilbert & Li, 2013; Herzog & Manassi, 2015; O’Callaghan, Kveraga, Shine, Adams & Bar, 2017; de Lange, Heilbron & Kok, 2018). By now, it is evident that predictions and context modulate the processing of sensory stimuli. Following the Bayesian accounts of perception (e.g. Friston 2005; 2010), higher level cortical structures generate predictions (priors) based on previous experience and beliefs. These priors are compared to lower level sensory information for mismatches (prediction error signals) which are used to update higher level representations when necessary (see also Clark, 2013; Hohwy, 2013). Such predictive coding models suggest that perception is always influenced by priors and that the extent of this influence depends on the relative precision attributed to priors and prediction errors. In general, taking priors into account when making sense of sensory information ensures that our perception of the world is fast and seamless, allowing for efficient behaviour and economical use of neurobiological resources, but it can also lead to perceptual errors or illusions when the incoming sensory information is deemed noisy or ambiguous. In certain instances when the expectation of a stimulus is very strong it can even result in the perception of something that is not actually present (e.g. Aru & Bachmann, 2017; Aru, Tulver & Bachmann, 2018; Powers, Mathys & Corlett, 2017).

According to the predictive coding framework, an imbalance between top-down signals and bottom-up sensory information is at the root of suboptimal perceptual processing and can be applied to explain a range of clinical symptoms (Fletcher & Frith, 2009; Adams, Stephen, Brown, Frith & Friston, 2013; Nour & Nour, 2015; Sterzer et al. 2018). For example, autism has been linked to weak or attenuated priors (Pellicano & Burr, 2012) and it has been suggested that in autistic people the balance between priors and bottom-up information processing is more skewed towards sensory information (Lawson, Rees & Friston, 2014; van de Cruys et al. 2014). On the other hand, certain positive symptoms of schizophrenia, such as hallucinations, are thought to be related to an increased use of prior knowledge or heightened priors (Powers et al 2017, Teufel et al 2015, Schmack et al. 2013), whereas delusions have instead been linked to weaker perceptual priors (for a review of the controversy see Sterzer et al. 2018). Some studies have reported that even healthy individuals with heightened levels of psychosis proneness display subtle perceptual alterations and a tendency to favour prior knowledge over incoming sensory evidence (Schwartzman et al. 2008; Teufel et al., 2015). Therefore, following a continuum hypothesis of psychosis which suggests that psychotic disorders are at the extreme end of a spectrum encompassing also the non-clinical population (e.g. Verdoux & van Os, 2002), it could be hypothesized that all individuals differ along a continuum of perceptual biases where some people rely more heavily on prior knowledge relative to sensory input than others, and vice versa.

### 1.3. Current study

The goal of this study was to investigate whether there is a general factor which one might call “relative reliance on priors” that would explain the behaviour of a given participant across several tasks. It is possible that there exist individual differences in how much participants *generally* rely on established prior knowledge and expectations compared to transient sensory information. This search for a general factor of reliance on priors is motivated by the literature where authors often make sweeping claims about priors (for example, suggesting that in autism the balance between priors and bottom-up information is tilted towards sensory information or that hallucinations are related to overweighting of priors). However, one could also consider that given a specific task, the brain makes use of specific priors that are activated by the particular stimuli and task structure. If the latter were true then any claim about “priors” in general would be misleading (for example, autistic individuals might have attenuated priors in some tasks, but not in others; people who hallucinate might have stronger priors in particular stimulus domains, but not in others). In this case priors never can and never should be dissociated from the tasks and stimuli with which they are studied. The debate on the validity of abstract optimal models vs task dependent suboptimal models of perception continues (e.g. Rahnev & Denison, 2018). In the present study we aimed to contribute to this by focusing on the universality of the effects of priors.

To distinguish between these ideas - a general process underlying all priors or the existence of many task-specific priors - we conducted an experiment where each participant was subject to four different tasks. A multi-paradigm approach allows us to explore how much individuals differ regarding the effects of priors on objective performance and subjective perceptual experience, and whether these differences are similar across various tasks. The tasks were selected on the basis of two main criteria. First, there exist consistent findings reporting a discrepancy between the subjective perceptual experience and objective characteristics of the task stimulus. Second, the respective discrepancy can be explained by priors overriding the present sensory information. Four visual perception paradigms were chosen: (1) perception of illusory contours, (2) blur detection, (3) Mooney face recognition, and (4) representational momentum effect.

The **illusory contours task** involves the subjective perceptual experience of contours in the absence of actual local border edges or lines (e.g. Kanizsa, 1976; Bachmann, 1978). In this instance, exemplified by the well-known Kanizsa figures or the corresponding Varin figures (Varin, 1971), previous cumulative experience with more typical shapes (e.g. squares and circles), as suggested by the specific arrangement of the inducing visual elements, creates the expectation that a square placed on top of four circles is a more likely interpretation than four symmetrically placed Pac-Man shapes.

A similar effect can be observed in a **blur detection task** recently introduced by Lupyan (2017). He demonstrated that people perceive meaningful words more sharply than their meaningless, nonword counterparts, although the letters in both strings remain the same and have identical spatial frequency content. Again, the effect presumably emerges as prior knowledge and experience with words create stronger priors for familiar letter strings compared to scrambled letter combinations and hence the subjective visual quality of intelligible words appears more detailed.

Another paradigm which was included in the study is the **Mooney face recognition task,** as Mooneys or other two-tone images are frequently used to demonstrate the effects of predictive perception (e.g. Loth, Gómez & Happé, 2010; Teufel, Dakin & Fletcher, 2018). Mooney faces (Mooney, 1957) are high-contrast black and white images generated from photographs of faces that are initially construed as ambiguous but have been shown to become disambiguated into coherent percepts after original versions of the photographs are introduced (e.g. Gorlin et al., 2012). We propose that prior knowledge regarding facial configurations, as well as strengthened priors resulting from familiarized targets facilitate the sensory processing of incomplete and noisy input leading to the images being identified as faces faster and more accurately. Additionally, a task specific expectation to perceive faces may lead to increased false positive rates in some, more so than other subjects.

Lastly, we included the **representational momentum task**. In this task, people observe a target moving smoothly along a straight trajectory (e.g. horizontal or vertical axis) until it disappears, typically without warning. A forward shifted mislocalization (difference between the perceived vanishing point and the actual vanishing point) is consistently found in analogous tasks (Freyd & Finke, 1984; for review, see Hubbard, 2005; 2018) and the effect has been shown to depend on the predictability of the direction of the target (Kerzel, 2002). The phenomenon of momentum in this instance captures the role of prediction (that the movement will continue in the expected direction), until the bottom-up correction (that the movement has stopped) reaches perception. In this manner, it would be possible to measure individual differences in relying on prediction, as a function of displacement size.

With this approach we can distinguish between several competing hypotheses. As already introduced above, one hypothesis posits that individuals are characterized by varying degrees of how much priors contribute to perception, and that these differences are systematic across separate perceptual tasks. More specifically, if there exists a common general factor for the expression of the effects of priors, people who experience illusory contours more clearly might also report larger displacement in the representational momentum task, benefit more from the introduction of priors in the Mooney task and perceive meaningful words relatively more sharply than meaningless words.

Alternatively, it is possible that individual differences in the proneness to rely on priors might not be expressed universally across all four tasks, but only within subsets of tasks, relating to more specific types of priors. For instance, Series and Seitz (2013) have suggested that visual expectations fall into two broad categories (structural and contextual) based on how the priors were acquired and the extent to which they can be generalized across different environments. Finally, it is also important to consider the hypothesis that priors are always related to specific stimuli and task demands and thus general claims about priors are misleading.

Questionnaires were added to the experimental setup to investigate how these individual differences in the effects of priors on perception relate to measures of the autism spectrum and schizotypal personality in the non-clinical population. Based on previous research we hypothesized that individual measures of the impact of priors on perception display a positive correlation with schizotypy scores (Teufel et al. 2015) and a negative correlation with autistic trait scores (Aru et al., 2018).

## 2. Materials and Methods

### 2.1. Participants

Forty-four subjects participated in the experiment (15 males, 29 females). The sample size was determined by practical constraints. Participant ages ranged from 19 to 43 years old (M = 28 years, SD = 4.9). All participants were healthy and had normal or corrected-to-normal vision. Five participants were left-handed. All participants gave written informed consent prior to participation.

The study was approved by the ethics committee of University of Tartu and the experiment was carried out in compliance with national legislation and the Declaration of Helsinki.

### 2.2. Procedure

Participants were seated in a dimly lit room, 90 cm from the monitor (SUN CM751U; 1024×768 pixels; 100 Hz refresh rate).

Each of the four experimental tasks was preceded by a short introduction and 8-10 practice trials during which participants had the chance to rehearse the upcoming behavioural task. The experiments were run on custom scripts programmed in Python. As the focus of the study was not on a comparison of tasks, but on the exploration of individual differences in the common effects across all paradigms, the experimental tasks were presented in a fixed order, as follows: 1. illusory contours, 2. blur detection, 3. Mooney face recognition, 4. representational momentum. See Figure 1 for illustrations of the task stimuli.

**Figure 1.**
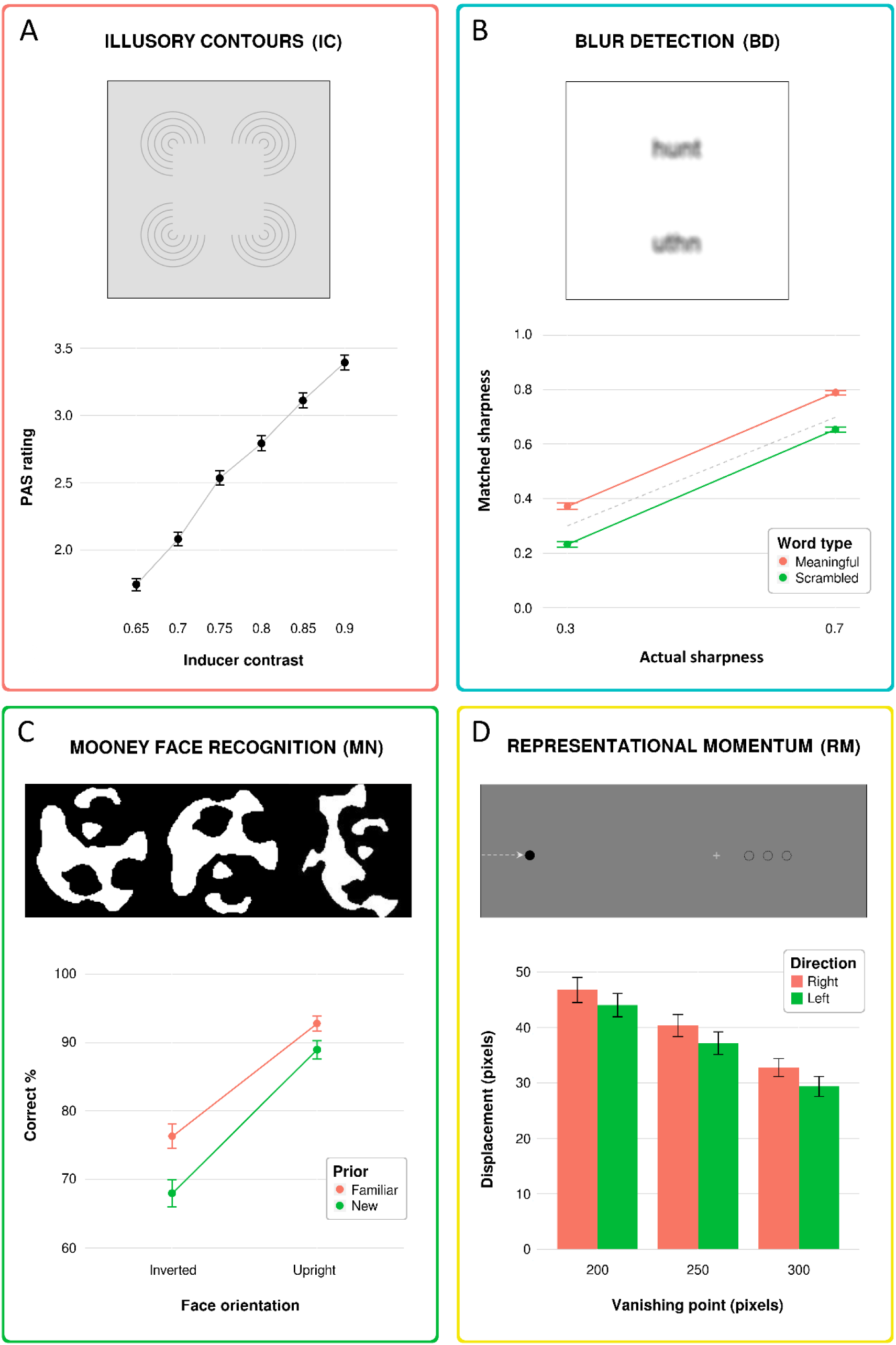
Illustrations of the four visual paradigms (upper panels) and their main behavioural effects including 95% confidence intervals (lower panels). A. Illusory contours (IC): example stimulus and mean PAS ratings at each level of inducer contrast. B. Blur detection (BD): meaningful (upper) and scrambled (lower) word pair at the same objective level of blur; actual sharpness of the standard word and mean adjusted sharpness of the target word in the meaningful (MN) and scrambled (SC) standard condition (greater values indicate higher levels of sharpness; note that in Lupyan’s original figure the scale is reversed). The grey dashed line represents veridical matched sharpness levels. C. Mooney face recognition (MN): from left to right – inverted Mooney face, upright Mooney face, and scrambled target; mean percentages of correctly recognized faces across conditions (not including trials with scrambled images). D. Representational momentum (RM): illustrative excerpt of the stimulus screen in the representational momentum task during a trial in which the stimulus (black circle) enters from the left side of the screen and vanishes in one of the three possible locations (depicted as empty circles for illustration purposes) on the right side of the screen. A light cross then appeared which participants could adjust to mark the apparent vanishing point (note that the depicted cross is larger and higher in contrast for the purpose of clarity); mean displacement sizes across conditions.

#### 2.2.1. Illusory contours task

The target was a five-line Varin figure consisting of four Pac-Man shaped inducers with a size of 2.2 degrees of visual angle (between 10 and 70 px pixels in radius) and a virtual square of 3.3 degrees of visual angle (side length 200 px). The contrast of the inducers varied across six different levels (65-90% between the lightest and darkest endpoint); targets were presented on a light grey screen (30.4 cd/m²) equal to 50% of nominal contrast. Inducers of every contrast level were each displayed 20 times in randomized order (120 trials in total).

Each trial began with the presentation of a fixation cross in the centre of the screen. The Varin figure was presented for 1500 ms, during which participants were able to view the figure without restrictions. The participants then had to indicate on a four-point perceptual awareness scale (PAS) (Overgaard, Rote, Mouridsen, & Ramsøy, 2006) how clearly they were able to perceive the subjective illusory contours of the square, using the keys A,D,J,L. The respective levels of the visibility scale were “no experience”, “a weak experience”, “an almost clear experience” and “a clear experience” of the contours (in the Estonian language). The four levels of the scale were displayed on a response screen at the end of every trial and participants were instructed in detail before the task on how to use the scale to describe their perceptual experience.

#### 2.2.2. Blur detection task

The blur detection task was a slightly modified version of the perceptual matching task described by Lupyan (2017).

The stimuli were 15 different four-letter words selected from a word frequency list of the most commonly encountered words in the Estonian written language, printed in black Arial font. An additional 8 words were selected for the practice trials. For each word a matched nonword was created by scrambling the letter order, while making sure that the chosen nonword did not resemble any other meaningful letter string. The words were then blurred using Adobe Photoshop version CC 2015 “Field Blur” filter. A full list of the words and nonwords used in the experiment can be found on the project OSF page (https://osf.io/mxdbv/).

On each trial, participants were presented with two blurred words on a white background (49.9 cd/m²). A meaningful word was always paired with its scrambled counterpart and vice versa. The stimuli were horizontally centred – the standard word was displayed at 90 pixels above the centre, and the target word at 90 pixels below the centre of the screen. Participants were instructed to adjust the target to the same level of blurriness as the standard by using the ‘up’ and ‘down’ arrow keys. The standard and target words were of the same size, subtending approximately 0.9 (height) and 2.5 (width) degrees of visual angle.

The standard stimuli were presented at two levels of blur, 0.3 and 0.7 (30% and 70% between the blurriest and sharpest endpoint) and the target was set to the midpoint at 0.5 level of blur, so that participants had to increase or decrease blurriness of the target on an equal number of trials. The experiment consisted of two blocks of 60 randomized trials, so that each individual string of letters was presented a total of 8 times. The second block was identical to the first, except that the positions of the standard and target on the screen were switched.

#### 2.2.3. Mooney face recognition task

Black-and-white Mooney images were created using photos of faces selected from the freely available “Labeled Faces in the Wild” (http://vis-www.cs.umass.edu/lfw/) database. Pictures were selected based on the unfamiliarity criterion (i.e. famous or otherwise easily recognizable people were excluded) and suitable lighting conditions for Mooney generation. The images were roughly the same size and luminosity. Upright faces, inverted faces and scrambled images were used as targets. Scrambled images were created by manually reconfiguring the elements of each Mooney face until it no longer resembled a face.

The Mooney faces task comprised of 10 experimental blocks of 32 trials, as well as an additional pre-block intended to familiarize the participants with the task and to provide them with prior exposure to images presented in the next block. Each block consisted of the already seen and previously unseen Mooney faces (20 trials), as well as scrambled Mooney images (12 trials). The size of the stimuli was approximately 8 degrees of visual angle and they were presented on a nominally black screen (0 cd/m²) for 10 ms. Participants were instructed to give a speeded response as to whether the image depicted a face (upright or inverted) or not (scrambled), by pressing the letter L or A, respectively. In between the blocks of Mooney targets, four original photos of people were shown. Participants were instructed to attend to the pictures and answer a few simple questions about the people depicted in the photos (“Was the person in the photo a man or a woman?”, “Was the person in the photo more or less than 50 years old?”). Mooney faces generated from these photos were used as the new condition in the preceding block and the familiar condition in the following block. As a control condition of the effect of strengthening priors due to the introduction photos, 20% of the Mooney trials included familiar and new Mooney images for which original photos were not shown. In other words, in this condition familiar Mooneys were only familiar due to repetition from the previous block.

#### 2.2.4. Representational momentum task

The target was a filled black circle, approximately 0.40 degrees of visual angle (12 pixels in radius), presented on a grey background (15.8 cd/m²). The circle entered from the midpoint of either the left or right edge of the screen and moved continuously, with invariant speed across the screen along a straight imaginary horizontal line. The target velocity was obtained by a shift of 6 pixels between successive images, resulting in a moderate velocity of approximately 600 px/s (equivalent of 10 deg/sec). Target characteristics and velocity were kept the same throughout the experiment, as they have been shown to affect illusory displacement size (Hubbard, 2005). The circle then vanished at one of three possible locations (200, 250, or 300 pixels from the centre of the screen) along the axis of motion. After the target vanished, a small inconspicuous white cross appeared at a randomized location within 40-50 pixels behind the spatial location of the vanishing point (in relation to the trajectory of movement, i.e. closer to the centre of the screen). These minor deviations in where the cross appeared were introduced to prevent the participant from using it as a constant reference point. The participants then indicated their perceived vanishing point of the target by positioning the cross over where they last saw the circle using the ‘left’ and ‘right’ arrow keys. Arrow keys were used instead of the mouse cursor so that participants would not be forced to look away from the vanishing point and potentially forget its exact location. This was achieved by participants holding their fingers on the respective keys throughout the experiment. Each participant received 90 trials in randomized order.

### 2.3. Questionnaires

After the experiment, all participants completed two questionnaires: the Autism Spectrum Quotient (AQ) questionnaire (Baron-Cohen et al., 2001) with 50 items and the Schizotypal Personality Questionnaire-Brief (SPQ-B) questionnaire (Raine & Benishay, 1995) with 22 items, both of which had previously been translated to Estonian, back-translated for validation and piloted.

### 2.4. Data preprocessing

In order to acquire measures that would be representative of each participant’s overall tendency to rely on priors in their perceptual responses, behavioural data was cleaned and individual scores were calculated for every paradigm in a task-specific manner, as described in the following sections.

All data preprocessing and the consequent statistical analyses were performed using R software (https://www.R-project.org/; version 3.3.0).

### 2.4.1. Illusory contours task

To calculate individual scores, a curve was fitted to every participant’s set of responses (PAS scores were coded as 0.25, 0.5, 0.75, 1) across six levels of inducer contrast using the *quickpsy* package in R. Individual thresholds of subjective visibility were extracted at 50% of the maximum rating to capture the threshold of inducer contrast where a participant was more likely to report having seen the illusory contours than not. The threshold measure is henceforth referred to as the “subjective vividness” score. In addition, the mean PAS rating was calculated for each participant.

### 2.4.2. Blur detection task

In the instructions prior to the experiment, participants were advised not to spend an inordinate amount of time on one word pair, and this was also noted during practice trials. However, there was no external limit to how much time was spent per trial. Reaction times were recorded, and these varied substantially among participants. To minimize the effects of alternating strategy on the results, trials where a participant’s reaction time was longer than 2.5 standard deviations of their individual mean were excluded. Altogether 2.77% of trials were removed as a result.

Response accuracy was measured as the difference between the adjusted sharpness of the target stimulus and the actual sharpness of the standard stimulus. Individual scores were calculated by subtracting a participant’s mean response accuracy in the scrambled word condition from the mean response accuracy in the meaningful word condition, resulting in the measure “benefit of meaning”. The larger the difference, the more of an effect the meaningfulness of the word had on blurriness evaluation response accuracy.

### 2.4.3. Mooney face recognition task

Trials where the reaction time was longer than 2.5 standard deviations above the individual mean were excluded from further analysis. As participants were instructed to give as accurate and speedy responses as possible, we can assume that very long reaction times were the result of distraction or other confounding condition not related to the main task. Thereby, 1.93% of trials were removed.

Three individual scores related to effects of priors were calculated for the Mooney task. First, the mean percentage of correctly recognized faces in the upright face condition was subtracted from the percentage obtained in the inverted face condition, referred to as the “benefit of orientation” score. Second, each participant’s false positive rate of face recognition was calculated as the ratio of scrambled targets mistaken for faces. This variable was transformed to a logarithmic scale due to a positive skew (skewness 1.92) and is referenced as the “false positive” score.

Lastly, the mean percentage of correctly recognized faces in the new face condition was subtracted from the percentage obtained in the familiar face condition, as a measure of the benefit resulting from a previously seen target (measure “benefit of prior”). However, the experimental manipulation check revealed no statistical difference in “benefit of prior” scores between the condition where the original photo was presented and the control condition where no photo was shown (t(43) = 1.62, p = .11), which implies that the difference between new and familiar faces was mainly due to target repetition, rather than the introduction of specific priors. As this measure also exhibited very low split-half reliability (−1.24), we removed this measure from the final analyses, leaving two separate individual measures for the Mooney task.

### 2.4.4. Representational momentum task

Displacement size was calculated as the difference in pixels between the judged vanishing point and the actual vanishing point of the target. The no-report trials where participants had not adjusted the cross at all were removed (49 trials, 1.2% of data). In addition, trials where the recorded response was not within 2.5 standard deviations of a participant’s mean displacement were excluded from further analysis. As a result, altogether 2.7% of data were rejected in the representational momentum task.

In order to account for the main effect of vanishing point while still retaining data from all three conditions, we used a linear mixed model with the *lmer* function of the *lme4* package. Displacement size was modelled as a function of vanishing point as the fixed effect (lmer(DP~VP + (1+VP|ID))). Individual random effect coefficients were extracted (using the *ranef* function) to illustrate the individual differences in displacement size which were not a result of the overarching main effect of vanishing point, thereby arriving at the measure “displacement”.

## 3. Results

### 3.1. Descriptive and inferential statistics for individual tasks

#### 3.1.1. Illusory contours task

As expected, illusory contours in the trials with darker inducers were rated higher on the subjective visibility scale (Figure 1A). Across the sample, higher contrast levels predicted higher values of the threshold measure of the PAS ratings. The linear regression equation proved significant (F(1,262) = 274.3, p < .0001), with an R^2^ of 0.51.

In addition to the threshold measure, the mean PAS rating was calculated for each participant (M = 2.61, SD = 0.51). Since the two individual measures were highly intercorrelated (Pearson’s r = -.96, p < .0001), only the threshold measure was included in the final analyses.

#### 3.1.2. Blur detection task

A repeated measures ANOVA with the factors word type (meaningful, scrambled) and blur level (0.3, 0.7) was conducted. As anticipated, there was a significant main effect of word type (F(1,43) = 157.99, p < .0001, η^2^_G_ = 0.490), but neither an effect of blur level (F(1,43) = 2.10, p = .154) nor an interaction (F(1,43) = 0.34, p = .563). Overall, the results confirmed previous findings obtained by Lupyan (2017) – participants tended to adjust target stimuli to blurrier levels when matching them to scrambled standard stimuli (mean difference = - 5.76%, SD = 4.1%) and sharper levels when matching them to meaningful standard stimuli (mean difference = 8.10%, SD = 6.2%) (Figure 1B).

#### 3.1.3. Mooney face recognition task

A repeated measures ANOVA with the factors orientation (upright, inverted) and prior (new, familiar) was conducted. The mean performance level of face recognition across sample was 84.89% (SD = 6.01%). There was a significant effect of orientation (F(1,43) = 147.92, p < .0001, η^2^_G_ = 0.439) and prior (F(1,43) = 112.84, p < .0001, η^2^_G_ = 0.076) on the mean percentage of correct answers. Upright faces were recognized more correctly (M = 90.84%, SD = 6.70%) than inverted faces (M = 72.14%, SD = 13.02%) and familiar faces were recognized more correctly (M = 84.55%, SD = 8.64%) than new faces (M = 78.50%, SD = 9.71%). There was also a significant interaction of the two factors (F(1,43) = 15.20, p < .001, η^2^_G_ = 0.011), as the benefit of a familiar face was greater when the faces were inverted (Figure 1C).

There was also a significant effect of orientation (F(1,43) = 167.65, p < .0001, η^2^_G_ = 0.098) and prior (F(1,43) = 7.76, p = .008, η^2^_G_ = 0.002) on reaction times. Upright faces were recognized faster (M = 0.57, SD = 0.18) than inverted faces (M = 0.63, SD = 0.21) and familiar faces were recognized faster (M = 0.59, SD = 0.20) than new faces (M = 0.60, SD = 0.19). There was no interaction effect of the two factors on reaction times (F(1,43) = 0.31, p = .582).

There was a subtle but significant speed accuracy trade-off between reaction times and accuracy (Pearson’s r = -.12, p < .0001). Sensitivity measures (d′) of participants ranged from 1.11 to 3.63.

#### 3.1.4. Representational momentum task

To ensure that there was no learning effect resulting from 90 trials that could have affected displacement size a paired t-test was applied between the first and second half of trials. No significant difference in displacement size was found (t(43) = 0.86, p = .393).

For testing the potential effects of conditions on the size of displacement, a repeated measures ANOVA with the factors direction (left-right, right-left) and vanishing point (200px, 250px, 300px) was run. There was no significant effect of direction on displacement size (F(1,43) = 2.21; p = .144) or a direction x vanishing point interaction (F(2,86) = 0.14, p = .866). However, there was a significant effect of vanishing point (F(2,86) = 59.42, p < .001, η^2^_G_ = .085). Displacement, on average across the sample, appeared larger when the target disappeared closer to the centre than when it disappeared closer to the edge of the screen (see Figure 1D for illustration). Post-hoc pairwise t-test revealed that all pairs were significantly different (all t(43) > 6.42, p < .0001).

#### 3.1.5. Questionnaires

The mean AQ score for 44 participants was 17.59 (SD = 5.68), with a range from 5 to 27 (the maximum possible score is 50). According to the authors of the questionnaire, a score of 32 can be considered the critical minimum suggestive of an autism-spectrum disorder (Baron-Cohen, 2001), therefore our sample was well within the non-clinical range. The mean AQ score for male participants (M = 18.60, SD = 5.67) was slightly higher than for female participants (M = 17.07, SD = 5.71), but the difference was not significant (t(28) = 0.85, p = .404).

For the SPQ-B, the SPQ-B total score as well as the score for the subfactor Cognitive-Perceptual Deficits was calculated. The mean SPQ-B total score for 44 participants was 8.07 (SD = 3.59), with a range from 1 to 16 (the maximum possible score is 22). It has been suggested that individuals with schizotypal personality disorder (DSM-IV) would likely score a 17 or higher on the SPQ-B (Raine, 1995).

### 3.2. Pairwise correlations

Spearman-Brown corrected split-half estimates were calculated for task performance and individual measures. All measures showed high reliability (>0.7) and can be found in Table 1. The histograms of the five individual measures are shown in Figure 2. For ease of comprehension, the performance of all measures was ordered from smallest effect of prior to largest, so the directions of some of the variables were reversed. To assess the relationship between the individual scores of the different tasks and between the individual scores and the questionnaire measures, we conducted correlation analyses. As we were mainly interested in whether individuals who relied more on priors in one task also ranked higher in other paradigms, Spearman’s rank order correlation was used. All analyses were also carried out using Pearson’s r and provided similar results. Spearman’s rho effect sizes are reported in Table 1. Reported p values were left unadjusted for multiple testing, as Bonferroni or other corrections would reduce the statistical power to discover correlations between the tasks that are crucial for answering the experimental question of this study (see also Grzeczkowski et al. (2017) for further justification under circumstances similar to ours).

**Table 1.**
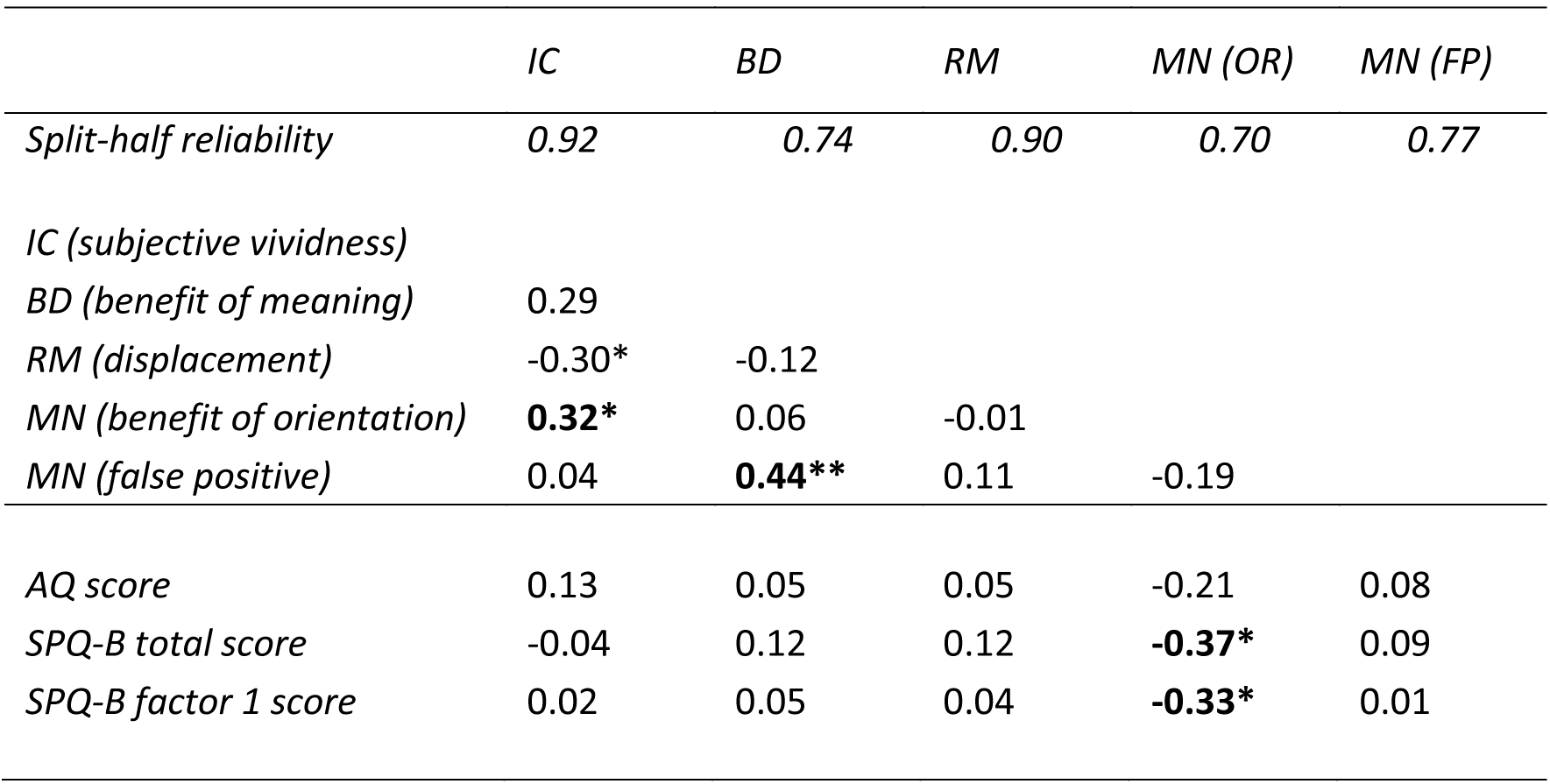
Spearman-Brown corrected split-half estimates of task reliability; Spearman’s correlations between the individual measures extracted from the behavioural tasks and questionnaires. N=44. P values unadjusted. * p < 0.05, ** p < 0.01.

**Figure 2.**
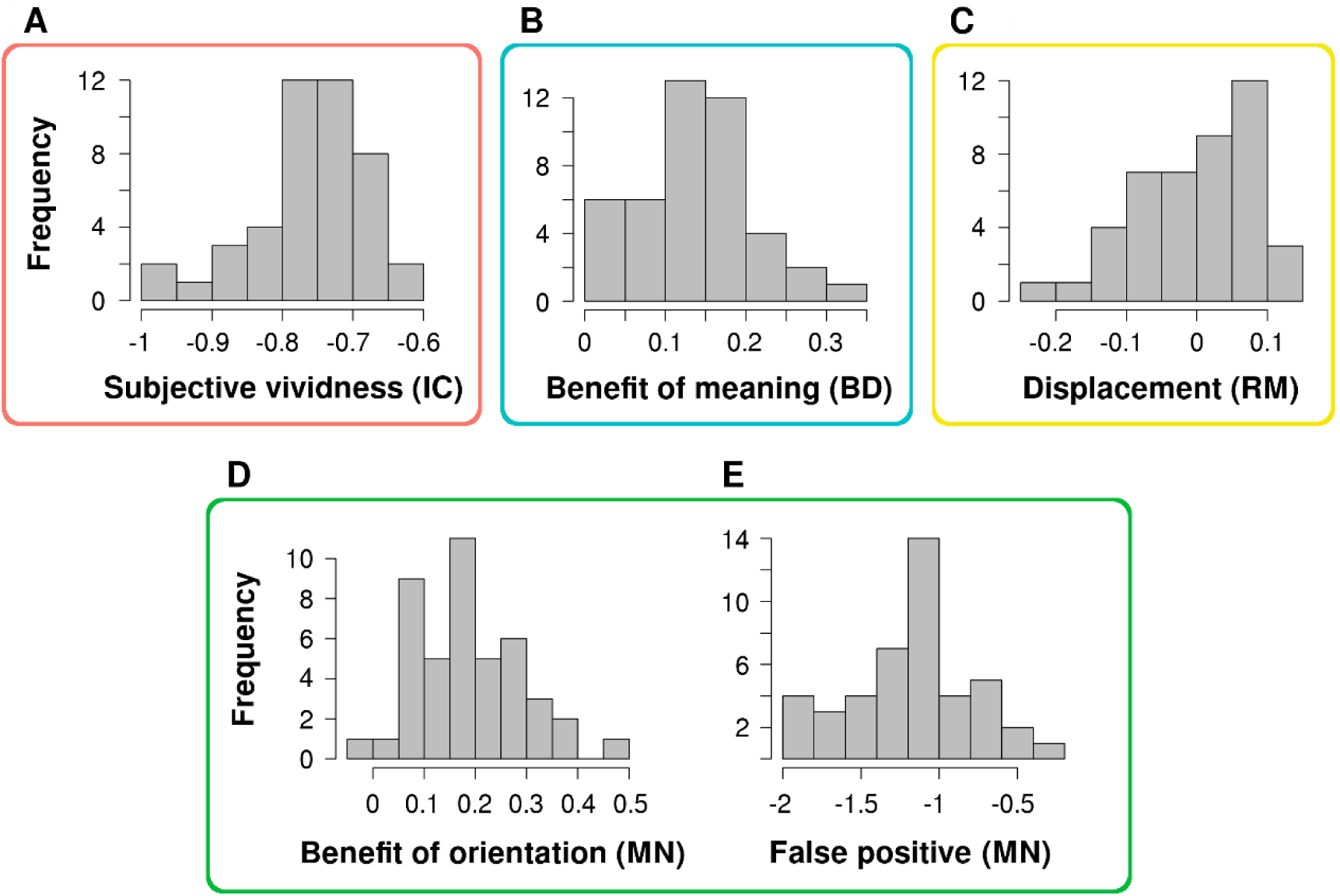
Histograms for the individual measures of the separate tasks: A. Illusory contours “subjective vividness” score; B. Blur detection “benefit of meaning” score; C. Representational momentum “displacement” score; D. Mooney “benefit of orientation” score; E. Mooney “false positive” score (logarithmic scale). It should be noted that the scores for IC and RM were reversed, therefore a higher score in all illustrated measures represents a larger effect of prior.

There was a significant positive correlation between the “subjective vividness” (IC) score and the “benefit of orientation” (MN) score (rho = .32, p = .034), and a negative correlation with the “displacement” (RM) score (rho = -.30, p = .048). “Subjective vividness” also exhibited a trend towards a correlation between the “subjective vividness” score and “benefit of meaning” (BD) score (rho = .29, p = .056). The “benefit of meaning” score was significantly correlated with the “false positive” (MN) score (rho = .44, p = .003).

The autism spectrum quotient did not exhibit any significant correlations with the five individual measures. However, there was an overall positive correlation between hit rates in the Mooney task and the AQ score (rho = .37, p = .013), including when only looking at responses to previously unseen Mooney images (rho = .39, p = .009).

The SPQ-B total score was negatively correlated with the “benefit of orientation” score (rho = -.37, p=.015). The Schizotypy subfactor of Cognitive-Perceptual Deficits also displayed a significant negative correlation with the “benefit of orientation” score (rho = -.33, p = .028), as well as the “displacement” score (rho = -.35, p = .020).

Nevertheless, it is important to note that none of the aforementioned correlations, except the correlation between the “false positive score” and “benefit of meaning” score, would survive a more conservative level of significance (p<0.01) or a correction for multiple comparisons.

### 3.3. Factor analysis

To explore the latent variable structure underlying the individual variation in the behavioural tasks, an exploratory factor analysis was performed on the five individual measures, using the maximum likelihood estimation, followed by Oblimin rotation and Kaiser normalization. Parallel analysis (Holm, 1965) was used to determine the number of factors to extract from the analysis. Two components were identified, altogether accounting for a cumulative 50% of the variance. To compare, a one-factor model would only explain 26% of variance. The first component explaining 25% of variance receives loadings (>0.3) from the “false positive” (MN), “benefit of meaning” (BD) and subjective vividness (IC) scores. The second component explaining an additional 25% of variance receives loadings from the “subjective vividness” (IC), “benefit of orientation” (MN) and displacement (RM) scores. Factor loadings for the two components are displayed in Table 2.

**Table 2.**
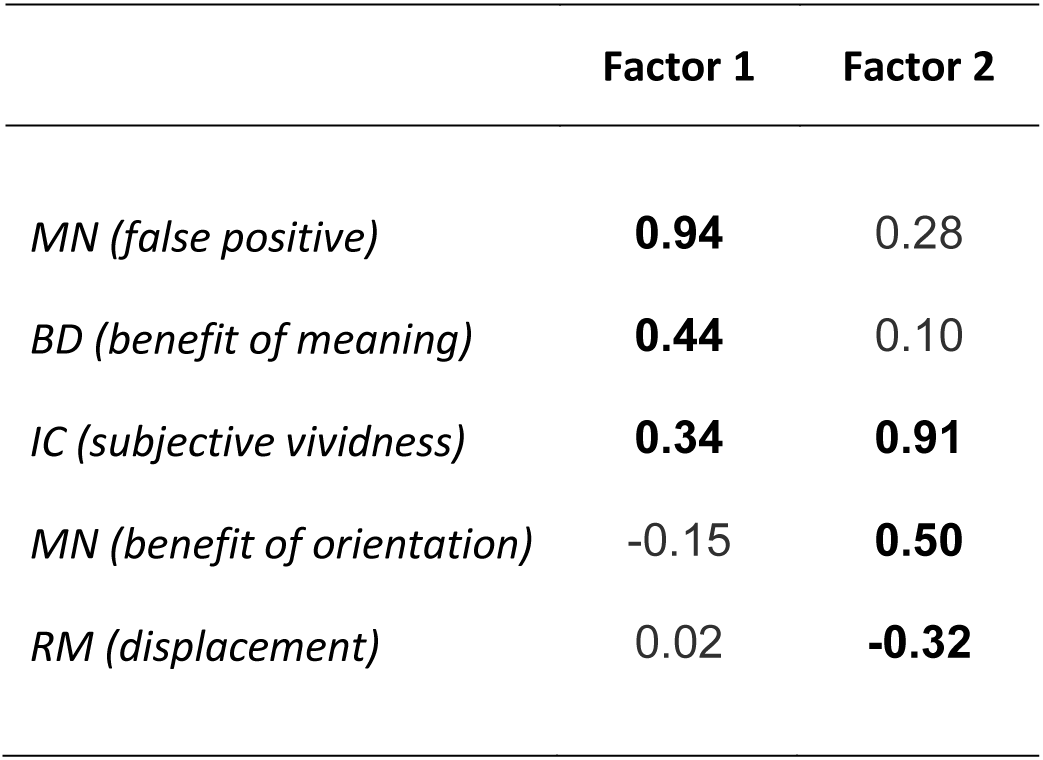
Factor loadings for the two rotated factors; loadings greater than 0.3 are highlighted in bold.

Although the factor analysis was exploratory, we conducted additional analyses to estimate the fit of the one factor model against the two factor model. The chi square goodness of fit statistic tests the hypothesis that the factor model fits the data perfectly. This hypothesis was rejected for the one-factor model (χ2(5) = 13.08, p = .023; RMSEA = 0.21), but not for the two-factor model (χ2(1) = 0.88, p = .349; RMSEA = 0). Even when removing the measure “displacement” (RM) from the analysis, which loaded most weakly to the two factors, the one factor model was not a good fit for the data (χ2(2) = 10.91, p = .004; RMSEA = 0.336) which provides further support to the claim that there is no general factor of relative reliance on priors.

## 4. Discussion

The quest for common factors of visual perception has so far led to mixed results, as the structure of individual differences in visual perception still remains unclear. Here we asked the question whether there exists a general factor for the relative reliance on priors when making sense of perceptually ambiguous or noisy stimuli. Participants completed four different behavioural tasks in which a discrepancy between objective sensory information and subjective perceptual experience has consistently been demonstrated. Performance on the tasks of each separate perceptual paradigm exhibited results consistent with previous findings. Taken as a whole, our results imply that one underlying factor does not explain the individual variance in the effects of priors on perception as measured by the four tasks. Rather, there seem to be different types of priors that are captured to a various extent by different tasks.

### 4.1. Priors affect perception in a variety of ways

The results from the correlational as well as factor analyses do not support the hypothesis for one common factor explaining the individual variance in the effects of priors on perception, such that would be analogous to the g factor considered in intelligence studies. Instead, our results based on the five extracted measures from four different experimental paradigms were best described by a two-factor structure, both factors explaining an equal portion of the cumulative variance (25%).

The two factors closely mirror the results from the pairwise correlation analyses and seem to relate to different levels of processing hierarchy. **The first factor**, which received the strongest loadings from the “false positive” (MN) score and the “benefit of meaning” (BD) score, can be interpreted as a factor reflecting higher level visual priors related to contextual expectations. In both tasks a high level prior with respect to the upcoming stimulus was evoked: in the Mooney task the participant was expecting to see faces, in the blur task the expectation was regarding letters and words. In both cases the prior is unspecific (face or word vs specific face or specific word), as participants pre-activated high-level generalized features relevant to the task. Hence, this type of prior could also be interpreted as an attentional effect steered by perceptual category information. **The second factor** loaded most strongly on the “subjective vividness” (IC) score and “benefit of orientation” (MN) score which are both related to the enhanced perception of familiar shapes compared to less likely interpretations of the image arrangement (i.e. a square overlaying four partly occluded discs instead of the four symmetrically arranged Pac-Man shapes, or an upright face shape instead of an inverted facial configuration), and thereby rely on more structural expectations acquired through life experience with basic shapes. In terms of processing hierarchy one could say that these measures capture mid-level priors. To a lesser degree, the “displacement” (RM) measure, as well as the “false positive” (MN) also loaded on this component. The main mechanism of the forward displacement effect in the representational momentum task also seems to make use of more hardwired mechanisms which have been shown to exist already early in life (Hubbard, Matzenbacher & Davis, 1999). Nevertheless, such a broad distinction of two categories of priors is probably too simplistic, since it only explains 50% of the variance. As the authors who proposed the categories of structural and contextual priors (Seriés and Seitz, 2013) emphasize, even structural priors can be modulated by context and shaped through new experiences in specific situations, resulting in more complex patterns of the effects of priors. Here we see that levels of processing hierarchy, as well as specific task demands are also likely to play a role.

In the current study a two-factor structure was able to explain the measures extracted from the four paradigms, which suggests that interactions with levels of processing hierarchy play an important role in accounting for the individual variance present in these tasks. It is important to note that we had four specific tasks and five specific measures. For these tasks and these measures the two-factor structure was the best approximation. However, if one were to employ more or simply different tasks, a different factor structure could appear. Hence, we are definitely not suggesting that all kinds of top-down effects on perception can be captured by these two factors. Rather, the bottom line of this research is that there is not a single common factor that reliably explains a substantial portion of the effects of priors on perception. Therefore, this study is in line with previous research that has reported low pairwise correlations between visual tasks and found either no common factors in visual perception (Cappe et al 2014; Goodbourn et al. 2012; Grzeczkowski et al 2017) or only few minor sub-factors of perception specific to a single task or narrow processing mechanism (Bosten et al. 2017; Verhallen et al. 2017; Ward et al. 2016). Likewise, the present study adds weight to arguments about the principal difficulty of establishing general prior-including models of optimal perceptual behaviour that could be applied to different tasks (Rahnev & Denison, 2018). Our work is consistent with the two remaining hypotheses: 1) that there are several different types of priors and 2) that priors are always dependent on the specific stimuli and task structure. In this work we could not conclusively distinguish between these two hypotheses, but our work together with previous literature suggests that there is no general factor of “relative reliance on priors”. Hence, when a given study observes some differences in the relative weighting of priors and sensory data between patient populations and healthy subjects then this finding should not be generalized to all tasks and stimuli – various priors have specific effects under particular stimulus circumstances.

### 4.2. Psychopathology

To a first approximation, autistic and schizotypal traits could be regarded as the opposite ends of imbalanced perceptual processing. Autism has been attributed to weak priors and an undue reliance on sensory information (e.g. Pellicano & Burr, 2012; van de Cruys et al. 2014; Lawson et al 2014), whereas the hallucinations and psychotic experiences characteristic of schizophrenia have been hypothesized to result from overly strong priors (e.g. Teufel et al, 2015; Powers et al. 2017; Schwartzman 2008; Schmack et al 2013). However, in this study we only found few sporadic links between individual differences in the effects of priors on perception and autistic or schizotypal traits.

The SPQ-B subfactor score was negatively correlated with the “displacement” (RM) score and the Mooney “benefit of orientation” score, indicating that participants with higher SPQ-B scores relied relatively *less* on prior knowledge in these two tasks. Although several studies have reported that schizotypal traits are related to heightened priors (Teufel et al. 2015, Powers et al 2017), the opposite has also been suggested. For example, Stuke and colleagues (2018) found that increased delusion proneness is instead associated with decreased use of priors in a perceptual task. They suggest that this distinction can be interpreted in terms of processing hierarchies, as low-level processing in schizotypal individuals has been associated with decreased priors, whereas higher level processing has been linked to increased use of priors (see also Schmack et al. 2013, Sterzer et al 2018). Indeed, both the representational momentum task, as well as the very basic processing of facial configurations can be construed as processes that take place in the lower hierarchical levels of the visual cortex.

The Autism Quotient score did not exhibit any correlations with the five individual measures, which falls in line with studies that have claimed that the use of priors is preserved in ASD (van de Cruys, Vanmarcke, van de Put & Wagemans, 2017). Nevertheless, there is still some controversy regarding the applicability of a trait questionnaire such as AQ as a proxy for ASD proper (Gregory & Plaisted-Grant, 2016), therefore caution should be applied when interpreting these results. It is also possible that the link between autism and use of prior knowledge is modulated by the volatility of the environment (Palmer et al., 2017; Lawson et al., 2017) or the granularity of priors (Kwisthout, Bekkering & Rooij, 2016) which we did not manipulate or control for in this experiment, which could also explain the lack of correlations with prior measures in the current study.

Overall, our results did not convincingly show that the autistic and schizotypal traits in the general population are reliably linked to individual differences in the effects of priors on perception, as measured by these tasks.

### 4.3. Future directions

Many questions regarding top-down influences on perception remain unanswered and the results across various studies often paint a muddled picture (de Lange, Heilbron & Kok, 2018). Here we emphasize the benefit of a multi-paradigm approach for studying individual differences in how priors affect perception. Previously, many different perceptual paradigms have been used to examine the effects of priors, but the demands of specific task structures can be multiple and varied, which makes it difficult to obtain robust and unambiguous results that could be generalized over multiple paradigms. In the current study we found that people who rely more on priors in one task may not necessarily exhibit the same tendencies in a different task. Some of the inconsistencies found in literature can therefore also be attributed to an overly generalized conceptualization of priors. A clarification of the nomenclature related to types of priors in various tasks is essential for acquiring a more comprehensive understanding of the diverse mechanisms underlying the effects of priors on perception.

The role of priors and prediction errors in autism and schizophrenia is still unclear partly due to the very complex symptomatology of psychopathology (Palmer et al., 2017; Sterzer et al., 2018). Research with non-clinical samples may prove useful in systematically studying the basic mechanisms underlying varying degrees of susceptibility to perceptual illusions (Stuke et al., 2018). However, evidence is still sparse and there have been conflicting findings regarding whether or not tendencies found in a clinical sample carry over to the general population (van de Cruys, Vanmarcke, van de Put & Wagemans, 2017; Williams, 2018; Karvelis, Seitz, Lawrie & Seriès, 2018). In this study we only looked at differences in the use of priors, but a more nuanced approach might be helpful. For example, it has been proposed that for a better understanding of perceptual biases, one needs to take into account the volatility of the environment (e.g. unpredictable changes in the stimulus-outcome relationship) (Palmer et al., 2017; Lawson et al., 2017; Schmack, Weilnhammer, Heinzle, Stephan & Sterzer, 2016), and that certain links with psychopathology may only emerge in an unstable environment (Cassidy et al. 2018). We note that in our present four experimental paradigms volatility was not manipulated and all tasks were non-volatile in the sense that the stimulus-outcome contingencies remained stable throughout the experiment. Future studies might benefit from taking this aspect into account. For instance, in the representational momentum task one could manipulate volatility by having blocks of trials where the moving stimulus disappears at a relatively common location (i.e. the vanishing point coordinate has low variance) and blocks where the stimulus disappears at various locations (i.e. the vanishing point coordinates have high variance). Another step of inquiry would be to validate this behavioural evidence with neural markers of visual processing to further clarify the variety of top down effects on perceptual experience.

### 4.4. Limitations

Due to the exploratory nature of our study, as well as a relatively small sample size, our results should mainly be viewed as a tentative step towards a better understanding of the individual differences in the effects of priors. More research is needed to clearly distinguish between the cognitive styles related to the involvement of different processing stages and different levels of predictive mechanisms. Our lack of significant pairwise correlations can partly be explained by low statistical power, as with a sample size of 44 subjects there is a power of 80% to detect effect sizes above 0.4, therefore future studies should examine whether more links between task measures could be detected with a larger sample size.

In the current study we employed four very different behavioural tasks where performance may have depended on the use of priors to a different degree. In principle it is even possible that there is one common prior, whose effect was confounded by differences between the tasks, rather than in the use of different priors. By only applying behavioural measures, the question of how to disentangle the contribution of priors from general task performance cannot fully be answered. However, based on general knowledge from the neuroscience of vision it is to be expected that different tasks engage different (types of) priors. For instance, the neural processing of faces happens along a separate pathway from processing words (Duchaine & Yovel, 2015; McCandliss, Cohen & Dehaene, 2003). While there may be some common aspect to how priors in both cases are applied, it is also clear that the strength of prior knowledge on these two pathways could be manipulated independently (e.g. by having much experience with faces but only little with words, or vice versa).

### 4.5. Conclusions

The present study aimed to investigate individual differences in the effects of priors on visual perception, using a multi-paradigm approach. Our results show that there is no single factor that would account for a substantial proportion of the individual differences in the effects of priors on perception, as captured by the four selected paradigms. Instead, we argue that perceptual priors likely originate from multiple separate sources and that the effects of priors on perception always depend on the specific stimuli and tasks. We find that with the current deluge of studies exploring top-down effects on perception, it is especially relevant to emphasize that experimental tasks should be selected more rigorously to accurately reflect the effects of priors on a more concrete level of processing and that results should be interpreted accordingly.

## Supplementary material

Complete data and analysis scripts are available on the project page on the Open Science Framework (https://osf.io/mxdbv/).

## Acknowledgments

We thank the reviewers Pieter Moors, Sander Van de Cruys and Michael Herzog for their helpful comments and suggestions.

This work is partly supported by Grants IUT20-40 and PUT1476 from the Estonian Ministry of Education and Research. KT was supported by the University of Tartu ASTRA Project PER ASPERA, financed by the European Regional Development Fund. JA was also supported by the European Union’s Horizon 2020 Research and Innovation Programme under the Marie Skłodowska-Curie grant agreement no. 799411.

